# Characterizing the pathogenic, genomic, and chemical traits of *Aspergillus fischeri*, a close relative of the major human fungal pathogen *Aspergillus fumigatus*

**DOI:** 10.1101/430728

**Authors:** Matthew E. Mead, Sonja L. Knowles, Huzefa A. Raja, Sarah R. Beattie, Caitlin H. Kowalski, Jacob L. Steenwyk, Lilian P. Silva, Jessica Chiaratto, Laure N.A. Ries, Gustavo H. Goldman, Robert A. Cramer, Nicholas H. Oberlies, Antonis Rokas

## Abstract

*Aspergillus fischeri* is closely related to *Aspergillus fumigatus*, the major cause of invasive mold infections. Even though *A. fischeri* is commonly found in diverse environments, including hospitals, it rarely causes invasive disease; why that is so is unclear. Comparison of *A. fischeri* and *A. fumigatus* for diverse pathogenic, genomic, and secondary metabolic traits revealed multiple differences for pathogenesis-related phenotypes, including that *A. fischeri* is less virulent than *A. fumigatus* in multiple animal models of disease, grows slower in low oxygen environments, and is more sensitive to oxidative stress. In contrast, the two species exhibit high genomic similarity; ~90% of the *A. fumigatus* proteome is conserved in *A. fischeri*, including 48/49 genes known to be involved in *A. fumigatus* virulence. However, only 10/33 *A. fumigatus* biosynthetic gene clusters (BGCs) likely involved in secondary metabolite production are conserved in *A. fischeri* and only 13/48 *A. fischeri* BGCs are conserved in *A. fumigatus*. Detailed chemical characterization of *A. fischeri* cultures grown on multiple substrates identified multiple secondary metabolites, including two new compounds and one never before isolated as a natural product. Interestingly, an *A. fischeri* deletion mutant of *laeA*, a master regulator of secondary metabolism, produced fewer secondary metabolites and in lower quantities, suggesting that regulation of secondary metabolism is at least partially conserved. These results suggest that the non-pathogenic *A. fischeri* possesses many of the genes important for *A. fumigatus* pathogenicity but is divergent with respect to its ability to thrive under host-relevant conditions and its secondary metabolism.

**Importance:** *Aspergillus fumigatus* is the primary cause of aspergillosis, a devastating ensemble of diseases associated with severe morbidity and mortality worldwide. *A. fischeri* is a close relative of *A. fumigatus*, but is not generally observed to cause human disease. To gain insights into the underlying causes of this remarkable difference in pathogenicity, we compared two representative strains (one from each species) for a range of host-relevant biological and chemical characteristics. We found that disease progression in multiple *A. fischeri* mouse models was slower and caused less mortality than *A. fumigatus*. The two species also exhibited different growth profiles when placed in a range of stress-inducing conditions encountered during infection, such as low levels of oxygen and the presence of reactive oxygen species-inducing agents. Interestingly, we also found that the vast majority of *A. fumigatus* genes known to be involved in virulence are conserved in *A. fischeri*, whereas the two species differ significantly in their secondary metabolic pathways. These similarities and differences that we identified are the first step toward understanding the evolutionary origin of a major fungal pathogen.

## Introduction

Aspergillosis is a major cause of human morbidity and mortality, resulting in over 200,000 life-threatening infections each year worldwide, and is primarily caused by the fungal pathogen *Aspergillus fumigatus* (1). Multiple virulence traits related to Invasive Aspergillosis (IA) are known for *A. fumigatus*, including thermotolerance, the ability to grow under low oxygen conditions, the ability to acquire micronutrients such as iron and zinc in limiting environments, and the ability to produce a diverse set of secondary metabolites (2).

*A. fumigatus* thermotolerance, a key trait for its survival inside mammalian hosts, is likely to have arisen through adaptation to the warm temperatures present in decaying compost piles, one of the organism’s ecological niches (3-5). The primary route of *A. fumigatus* colonization and infection is through the lung, where oxygen levels have been observed to be as low as 2/3 of atmospheric pressure, and a successful response to this hypoxic environment is required for pathogenesis (6, 7). *A. fumigatus* produces a diverse set of small, bioactive molecules, known as secondary metabolites, which are biosynthesized in pathways that exist outside of primary metabolism. Some of these secondary metabolites and their regulators have been shown to be required for severe disease in mouse models (8-10). Furthermore, a master regulator of secondary metabolism, *laeA*, is also required for full virulence in IA mouse model studies (11, 12).

Other species closely related to *A. fumigatus* are also capable of causing disease, but they are rarely observed in the clinic (2, 13-15). For example, *A. fischeri* is the closest evolutionary relative to *A. fumigatus* for which a genome has been sequenced (16, 17), but is only rarely reported to cause human disease (2). Recent evolutionary genomic analyses suggest that *A. fischeri* and *A. fumigatus* last shared a common ancestor approximately 4 million years ago (95% credible interval: 2 – 7 million years ago) (17). Why *A. fischeri*-mediated disease is less common than *A. fumigatus*-mediated disease remains an open question. Non-mutually exclusive possibilities include differences in ecological abundance, lack of species level diagnosis in the clinic of disease-causing strains, and innate differences in pathogenicity and virulence between the two species.

Previous studies have suggested that the difference in the frequencies with which the two species cause disease is unlikely to be solely due to ecological factors, as both can be isolated from a variety of locales, including soils, fruits, and hospitals (18-20). For example, approximately 2% of the fungi isolated from the respiratory intensive care unit at Beijing Hospital were *A. fischeri* compared to approximately 23% of fungal species identified as *A. fumigatus* (20). While *A. fischeri* is easily isolated from a variety of environments, only a few cases of human infections have been reported (21-24). Furthermore, numerous recent epidemiological studies from multiple countries that used state-of-the-art molecular typing methods were able to identify several rarely isolated pathogenic species closely related to *A. fumigatus*, such as *A. lentulus* and *A. udagawae*, as the source of 10-15% of human infections but did not identify *A. fischeri* in any patient sample (13, 14, 25-27).

If ecological factors and lack of precision in species identification cannot explain why *A. fischeri* is non-pathogenic and *A. fumigatus* is pathogenic, other factors must be responsible. An early genomic comparison between *A. fumigatus*, *A. fischeri*, as well as the more distantly related *Aspergillus clavatus* identified 818 genes that were *A. fumigatus*-specific (16). These genes were enriched for functions associated with carbohydrate transport and catabolism, secondary metabolite biosynthesis, and detoxification (16), raising the possibility that the observed differences in pathogenicity observed between *A. fischeri* and *A. fumigatus* have a molecular basis.

To gain further insight into why *A. fischeri*-mediated disease is less abundant than *A. fumigatus*-mediated disease, we took a multi-pronged approach to investigate phenotypic, genomic, and chemical differences between *A. fischeri* strain NRRL 181 and *A. fumigatus* strain CEA10. We observed that while *A. fischeri* is able to cause fatal disease in multiple animal models, its disease progression and response to multiple host-relevant stresses is markedly different than that of *A. fumigatus*. We also found that while the two organisms’ genomes are in general very similar, the sets of secondary metabolite pathways in each of them exhibit a surprisingly low level of overlap. Examination of the secondary metabolite profile of *A. fischeri* identified both previously isolated as well as novel compounds. Finally, construction of a mutant *A. fischeri* strain that lacked the *laeA* gene, a master regulator of secondary metabolism, and examination of its chemical profile suggested that LaeA-mediated regulation of secondary metabolism in *A. fischeri* closely resembles that of *A. fumigatus*. These results begin to reveal the molecular differences between *A. fischeri* and *A. fumigatus* related to fungal pathogenesis and suggest that a functional evolutionary genomic comparison between pathogenic and non-pathogenic species closely related to *A. fumigatus* harbors great promise for generating insights into the evolution of fungal disease.

## Results

### *A. fischeri* is significantly less virulent than *A. fumigatus* in multiple animal models of Invasive Pulmonary Aspergillosis (IPA)

In contrast to *A. fumigatus*-mediated disease, only a handful of cases of invasive fungal infections have been reported to be caused by *A. fischeri* (21-24). Given this contrast, we utilized two immunologically distinct murine IPA models to assess differences in pathogenicity and virulence between the two species. In a leukopenic murine model, *A. fischeri* NRRL 181 is significantly less virulent than *A. fumigatus* CEA10, in a dose dependent manner (Fig. 1). Using an inoculum of 1 × 10^5^ conidia, *A. fischeri* is completely attenuated in virulence, with 100% murine survival by day 15 post-fungal challenge. In contrast, inoculation with *A. fumigatus* results in 100% murine mortality by day 15 (Fig. 1A). Using a higher dose (2 × 10^6^) of conidia, both strains cause 90% mortality by day 14; however, the disease progression is markedly different. 80% of mice inoculated with *A. fumigatus* succumb to infection by day 4, whereas for mice inoculated with *A. fischeri*, the first mortality event occurs on day 5, and then one or two mice succumb each day until day 14 (Fig. 1B). Thus, despite the similar overall mortality at higher fungal challenge doses, *A. fischeri* is significantly less virulent than A. fumigatus in a leukopenic murine IPA model.

**Figure 1:**
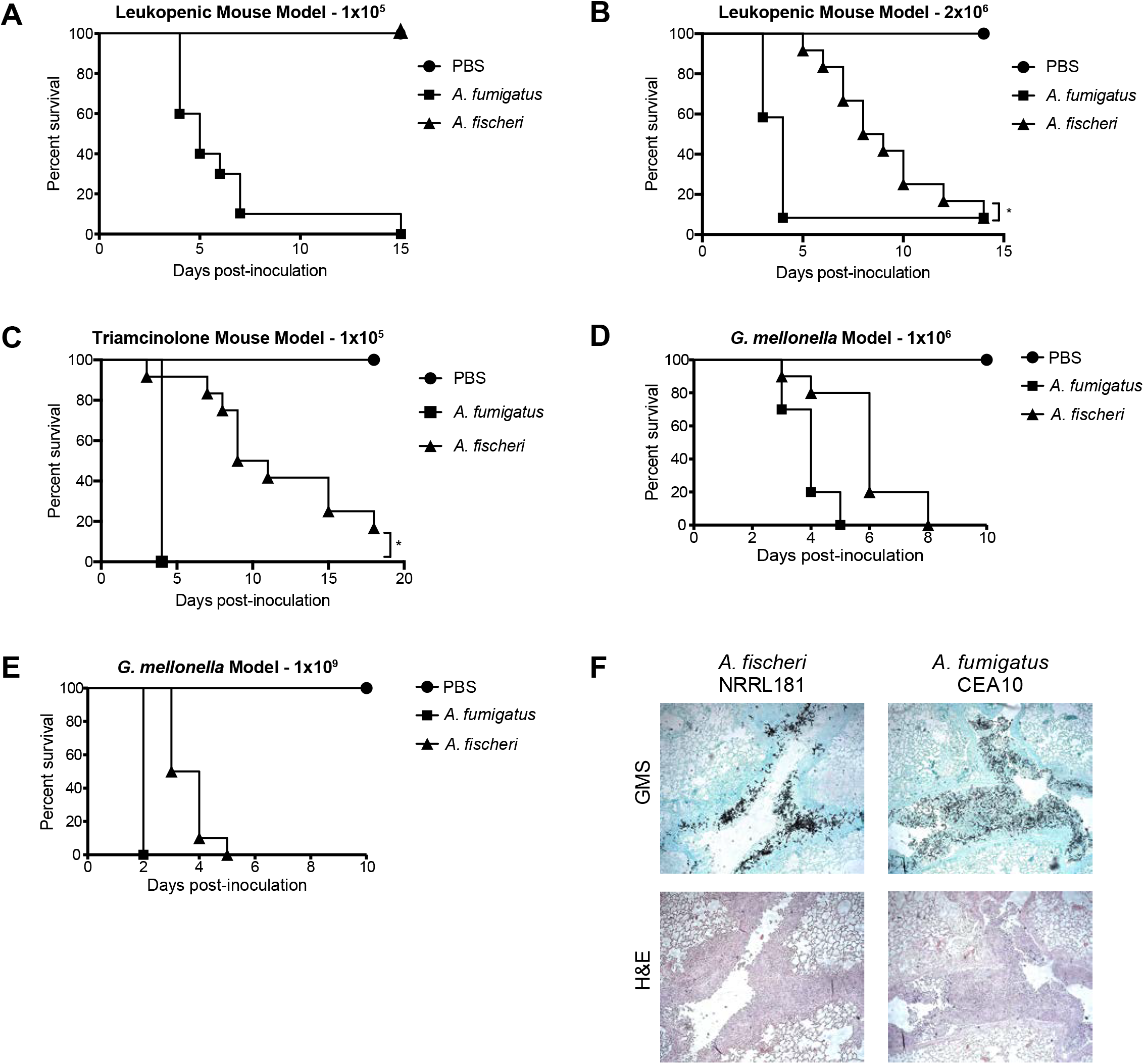
*A. fischeri* is significantly less virulent than *A. fumigatus* in multiple murine models of invasive pulmonary aspergillosis. AB) Cumulative survival of mice inoculated with 1 × 10^5^ (A) or 2 × 10^6^ (B) conidia in a leukopenic model of IPA. A) n=10/group B) n=12/group, 4/PBS. *p=0.0098 by Log-Rank test, p=0.0002 by Gehan-Breslow-Wilcoxon test. C) Cumulative survival of mice inoculated with 2e6 conidia in a triamcinolone model of IPA. n=12/group, 4/PBS. *p=<0.0001 by Log-Rank and Gehan-Breslow-Wilcoxon tests. D) and E) Cumulative survival of *G. mellonella* larvae inoculated with 1 × 10^6^ (D) or 1 × 10^9^ (E) conidia. 10 larvae were used per condition in all assays. Survival curves for *A. fischeri* and *A. fumigatus* were significantly different (p<0.003) in both Log-Rank and Gehan-Breslow-Wilcoxon tests for both inoculums. F) Histological sections from 3 days post inoculation in a triamcinolone model of IPA stained with H&E and GMS. Images were acquired at 100x.

As the patient population at risk for IA continues to change (28), we also tested a non-leukopenic triamcinolone (steroid)-induced immune suppression model and observed a significant reduction in virulence of *A. fischeri* compared to *A. fumigatus* (p<0.0001 by Log-Rank and Gehan-Breslow-Wilcoxon tests). All mice inoculated with *A. fumigatus* succumbed to infection by day 3; however, similar to the leukopenic model, mice inoculated with *A. fischeri* had slower disease progression as monitored by Kaplan-Meier analyses (Fig. 1C).

We observed similar pathogenicity and virulence results when using the *Galleria mellonella* insect larvae model of aspergillosis (Fig. 1DE). Both low (1 × 10^6^ conidia) and high (1 × 10^9^ conidia) inoculum experiments showed significant differences between the disease progression of *A. fischeri* (slower) and *A. fumigatus* (faster) in this insect model of fungal pathogenicity.

To better understand what is happening *in vivo* during disease progression with *A. fischeri* and *A. fumigatus*, histological analyses on lungs from the triamcinolone model 3 days post inoculation were utilized. Histological sections were stained with Gomori methenamine silver (GMS) to visualize fungal burden and with hematoxylin and eosin (H&E) stain to visualize host related pathology (Fig. 1F). Overall, mice inoculated with *A. fischeri* had similar numbers of fungal lesions as those inoculated with *A. fumigatu*s but the lesions caused by the two species were phenotypically distinct (Fig. 1F). In larger terminal bronchioles infected with *A. fumigatus*, there was greater fungal growth per lesion, and the growth was observed throughout the bronchiole itself, extending well into the lumen. These lesions are accompanied by substantial granulocytic inflammation and obstructs the airways surrounding the hyphae (Fig. 1F). In the lesions containing *A. fischeri*, the fungal growth is contained to the epithelial lining of the bronchioles. This pattern of growth is accompanied by inflammation at the airway epithelia, leaving the airway lumen largely unobstructed (Fig. 1F). The lack of airway obstruction during *A. fischeri* infection may contribute to the reduced virulence compared to *A. fumigatus*.

Although the distribution of the fungal lesions varies, there is still significant fungal growth in mice infected with *A. fischeri*, suggesting that *A. fischeri* is capable of growing within the immune compromised host. Indeed, we tested the growth rate of *A. fischeri* and *A. fumigatus* in lung homogenate as a proxy for growth capability within the nutrient environment of the host and observed no difference between the two strains (Fig. S1). These experiments show that in multiple models of fungal disease, *A. fischeri* is less virulent than *A. fumigatus*, although *A. fischeri* is still capable of causing disease using a higher dose and importantly, is able to grow within the immune compromised murine lung.

### When compared to *A. fumigatus*, *A. fischeri* differs in its response to several host-relevant stresses

Our *in vivo* experiments suggested that the lower virulence of *A. fischeri* is not solely a result of its inability to grow within the host. Therefore, we hypothesized that an additional contributing factor was the inability of *A. fischeri* to mitigate host stress. Nutrient fluctuation is a stress encountered *in vivo* during *A. fumigatus* infection (29). To assess differences in metabolic plasticity between the two species, we measured the two organisms’ growth on media supplemented with glucose, fatty acids (Tween-80), or casamino acids. Because low oxygen tension is a significant stress encountered during infection (6), and recently, fitness in low oxygen has been correlated to virulence of *A. fumigatus* (30), we measured growth of both species at 37°C in both normoxic (ambient air) and hypoxia-inducing (0.2% O_2_, 5% CO_2_) conditions. In normoxia with glucose, fatty acids (Tween-80), or casamino acids supplied as the carbon source, radial growth of *A. fischeri* was lower than that of *A. fumigatus* (Fig. 2). However, on rich media both organisms grew equally well (Fig. 2). We also observed a slower growth rate of *A. fischeri* compared to *A. fumigatus* in the first 16 hours of culture in liquid media supplied with glucose at 37°C. At 30°C, *A. fischeri* grew the same as, or better than, *A. fumigatus* except on Tween-80 where *A. fumigatus* had a slight advantage (Fig. S2). Also, *A. fischeri* grew substantially worse than *A. fumigatus* when grown at 44°C (Fig. S3). To determine relative fitness in hypoxic liquid environments, we measured the ratio of biomass in liquid culture in ambient air (normoxia) versus hypoxic (0.2% O_2_, 5%CO_2_) conditions. *A. fischeri* showed significantly lower fitness in hypoxic conditions, with about an 8.5-fold lower biomass than *A. fumigatus* (Fig. 3A). These data suggest that *A. fischeri* is less fit than *A. fumigatus* at 37°C and in low oxygen conditions, both of which have been shown to impact fungal virulence.

**Figure 2:**
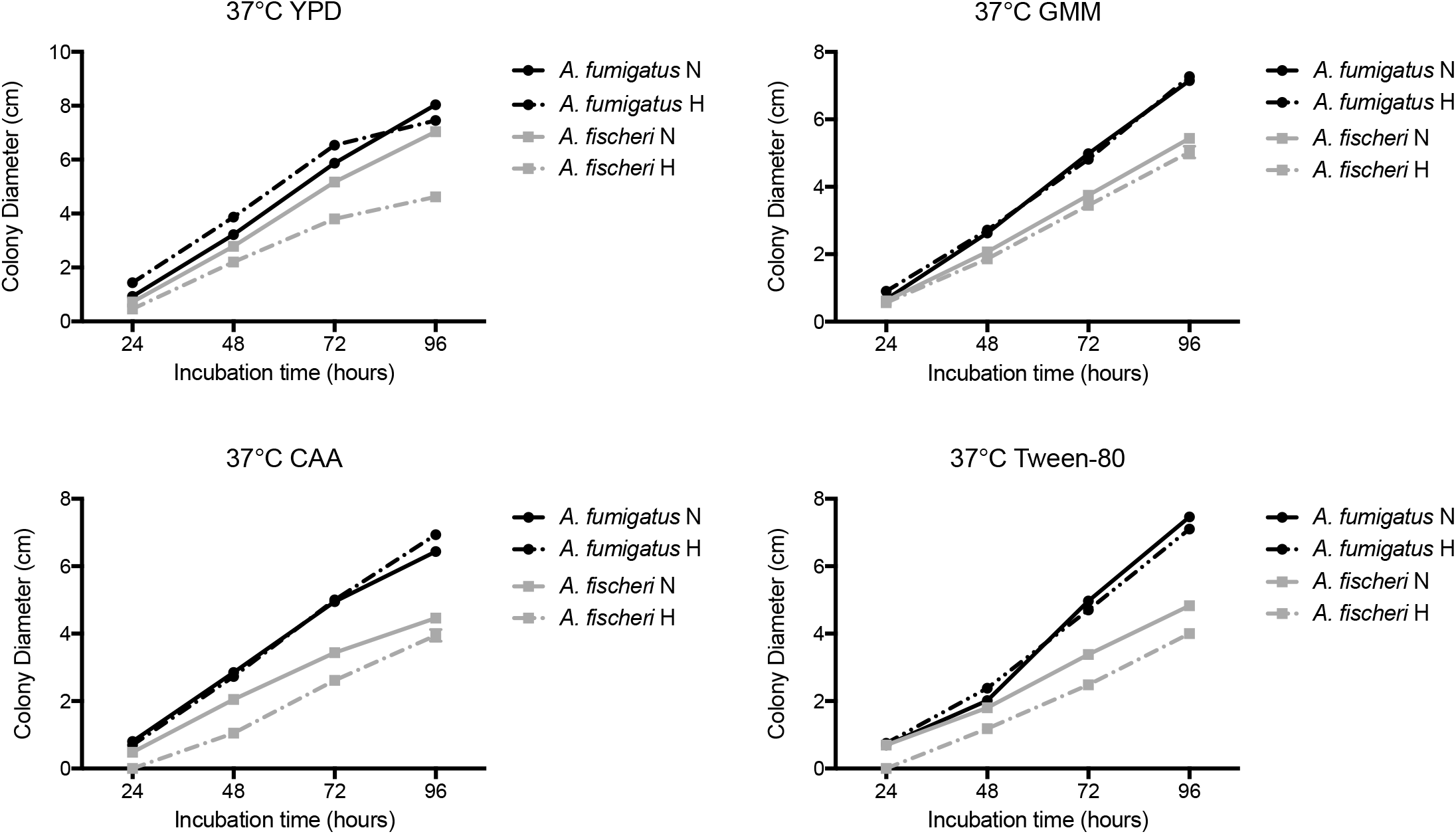
*A. fischeri* is unable to thrive under suboptimal metabolic conditions at 37°C. 1 × 10^3^ conidia were point inoculated on each plate then plates were incubated at 37°C in normoxia (N; ~21% oxygen, 5%CO_2_) or hypoxia (H; 0.2% O_2_, 5%CO_2_); colony diameter was measured every 24 hours. Mean and SEM of triplicates. CAA – Casamino acids; GMM – glucose minimal media.

**Figure 3:**
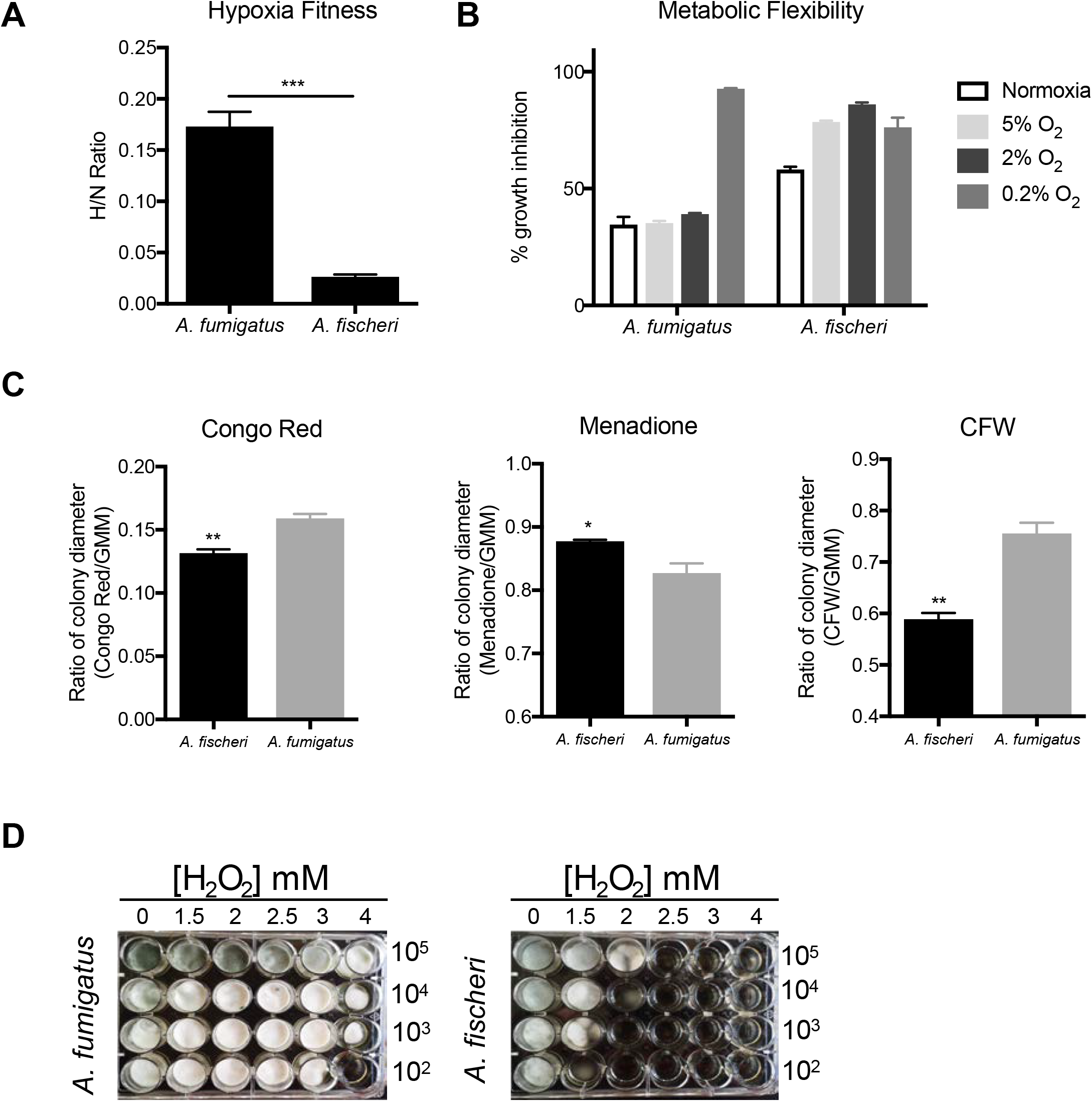
*A. fischeri* is more susceptible to multiple host-relevant stresses than *A. fumigatus*. A) Fitness ratio of *A. fumigatus* or *A. fischeri* during hypoxic vs normoxic growth (measured as the dry weight of cultures). Data represent mean and SEM of biological triplicates; ***p=0.0006 by Student’s t-test. B) Growth inhibition of strains grown on 1% lactate minimal media with 0.1% 2-deoxyglucose (2-DG) under a range of low oxygen conditions. C) *A. fumigatus* and *A. fischeri* were grown in the presence of the cell wall perturbing agent Congo Red (0.5mg/mL), the oxidative stressor Menadione (20 μM), or the chitin perturbing agent calcofluor white (CFW, 25μg/mL). Plates were grown for 96 hours at 37°C and 5% CO_2_. For all plates except Congo Red and its GMM control, 1e3 spores were plated. For Congo red and the control GMM plate 1e5 spores were plated. Student’s t-test was performed where *: p<0.05, **: p<0.01. D) Strains were grown for 48 h at 37°C in liquid complete medium supplemented with increasing concentrations of hydrogen peroxide.

Metabolic flexibility, or the ability for an organism to utilize multiple carbon sources simultaneously, has been suggested to provide a fitness advantage to *Candida albicans* during *in vivo* growth (31). Metabolic flexibility can be characterized using the glucose analog, 2-deoxyglucose (2-DG), in combination with an alternative carbon source available *in vivo*, such as lactate. 2-DG triggers carbon catabolite repression, which shuts down alternative carbon utilization pathways. However, in *C. albicans* this shut down is delayed and growth occurs on lactate with 2-DG (31, 32). We tested the metabolic flexibility of both *A. fumigatus* and *A. fischeri* and observed that while both species can grow in the presence of 2-DG on lactate, the growth inhibition of *A. fischeri* is higher (~60%) than that of *A. fumigatus* (~35%; Fig. 3B). Even under low oxygen conditions (5% and 2%), *A. fumigatus* maintains this metabolic flexibility except under extremely low oxygen conditions (0.2%), whereas *A. fischeri* shows even greater inhibition at all oxygen tensions of 5% or below. These data suggest that while both species exhibit some level of metabolic flexibility, *A. fumigatus* appears more metabolically flexible under a wider range of conditions than *A. fischeri*.

Next, we measured the susceptibility of *A. fischeri* to oxidative stress, cell wall stress, and antifungal drugs. Interestingly, we observed that *A. fischeri* is more resistant to the intracellular oxidative stress agent menadione than *A. fumigatus* but more susceptible to the external oxidative stress agent hydrogen peroxide (Fig. 3CD). As the *in vivo* levels of inflammation caused by the two species appeared different, we indirectly tested for differences in cell wall pathogen-associated molecular patterns using the cell wall perturbing agents Congo Red and Calcofluor White. *A. fumigatus* was significantly more resistant to both agents than *A. fischeri* (Fig. 3C), suggesting differences in the response to cell wall stress or in the composition and organization of the cell wall between the two species. These differences are likely important for host immune cell recognition and interaction, which in turn influences pathology and disease outcome.

Lastly, *A. fischeri* showed enhanced resistance relative to *A. fumigatus* for three of the four antifungal drugs tested (Table 1), consistent with previous experiments (33). Overall, our phenotypic data show that the response of *A. fischeri* to host-related stresses and antifungals is substantially different from that of *A. fumigatus*. Furthermore, our results suggest that increased growth capability of *A. fumigatus* in low oxygen and in high temperatures are two important attributes that likely contribute to its pathogenic potential compared to *A. fischeri*.

**Table 1.** *A. fischeri* shows enhanced resistance relative to *A. fumigatus* for several antifungal drugs.

| Strain | Posaconazole<br>[μg/ml] | Voriconazole<br>[μg/ml] | Itraconazole<br>[μg/ml] | Caspofungin<br>[μg/ml] |
| --- | --- | --- | --- | --- |
| *A. fumigatus* | 0.7 | 0.8 | 5 | 0.09 |
| *A. fischeri* | 2.4 | > 4 | > 24 | 0.06 |

### The proteomes of *A. fumigatus* and *A. fischeri* are highly similar, but their secondary metabolic pathways show substantial divergence

The large differences in virulence and virulence-related traits we observed between *A. fumigatus* and *A. fischeri* led us to investigate the genotypic differences that could be responsible. To describe the genomic similarities and differences between *A. fumigatus* and *A. fischeri*, we determined how many orthologous proteins and how many species-specific proteins were present in each genome using a Reciprocal Best BLAST Hit approach (34). We identified 8,737 proteins as being shared between the two species (Fig. S4), representing 88% and 84% of the *A. fumigatus* and *A. fischeri* proteomes, respectively, and 1,684 *A. fischeri*-specific proteins (16% of its proteome) and 1,189 *A. fumigatus*-specific proteins (12% of its proteome). To narrow our search for genes that are absent in *A. fischeri* but are important for *A. fumigatus* disease, we compiled a list of 49 *A. fumigatus* genes considered to be involved in virulence (Table S1) based on two previously published articles (35, 36) and extensive literature searches of our own. We observed that all but one of these virulence-associated genes were also present in *A. fischeri*, a surprising finding considering the substantial differences observed between the two species in our animal models of infection. The virulence-associated gene not present in *A. fischeri* is *pesL* (<u>Afu6g12050</u>), a non-ribosomal peptide synthase that is essential for the synthesis of the secondary metabolite fumigaclavine C and required for virulence in the *Galleria* model of *A. fumigatus* infection (37).

Since the only previously described *A. fumigatus* virulence-associated gene not present in the *A. fischeri* genome (i.e. *pesL*) is involved in secondary metabolism, we investigated the differences between the repertoire of secondary metabolic pathways present in *A. fumigatus* and *A. fischeri*. Using the program antiSMASH (38), we identified 598 secondary metabolic cluster genes distributed amongst 33 clusters in *A. fumigatus* (Table S2) and 786 secondary metabolite cluster genes spread out over 48 clusters in *A. fischeri* (Table S3). Of these 598 *A. fumigatus* genes, 407 (68%) had an orthologous gene that was part of an *A. fischeri* secondary metabolic gene cluster. This level of conservation of secondary metabolic cluster genes (68%) is much lower than the amount of conservation observed for the rest of the proteome (88%), illustrating the rapid rate at which fungal metabolic pathways evolve (39, 40).

We next directly compared the secondary metabolic gene clusters of the two organisms. An *A. fumigatus* gene cluster was considered conserved in *A. fischeri* if ≥ 90% of its genes were also present in an *A. fischeri* gene cluster and vice versa. We found that only 10 / 33 *A. fumigatus* gene clusters are conserved in *A. fischeri* and only 13 / 48 *A. fischeri* gene clusters are conserved in *A. fumigatus* (Fig. 4), a finding consistent with the low conservation of individual secondary metabolic genes between the two species. While only 10 *A. fumigatus* gene clusters were conserved in *A. fischeri*, many other clusters contained one or more orthologs of genes in *A. fischeri* secondary metabolic gene clusters.

**Figure 4:**
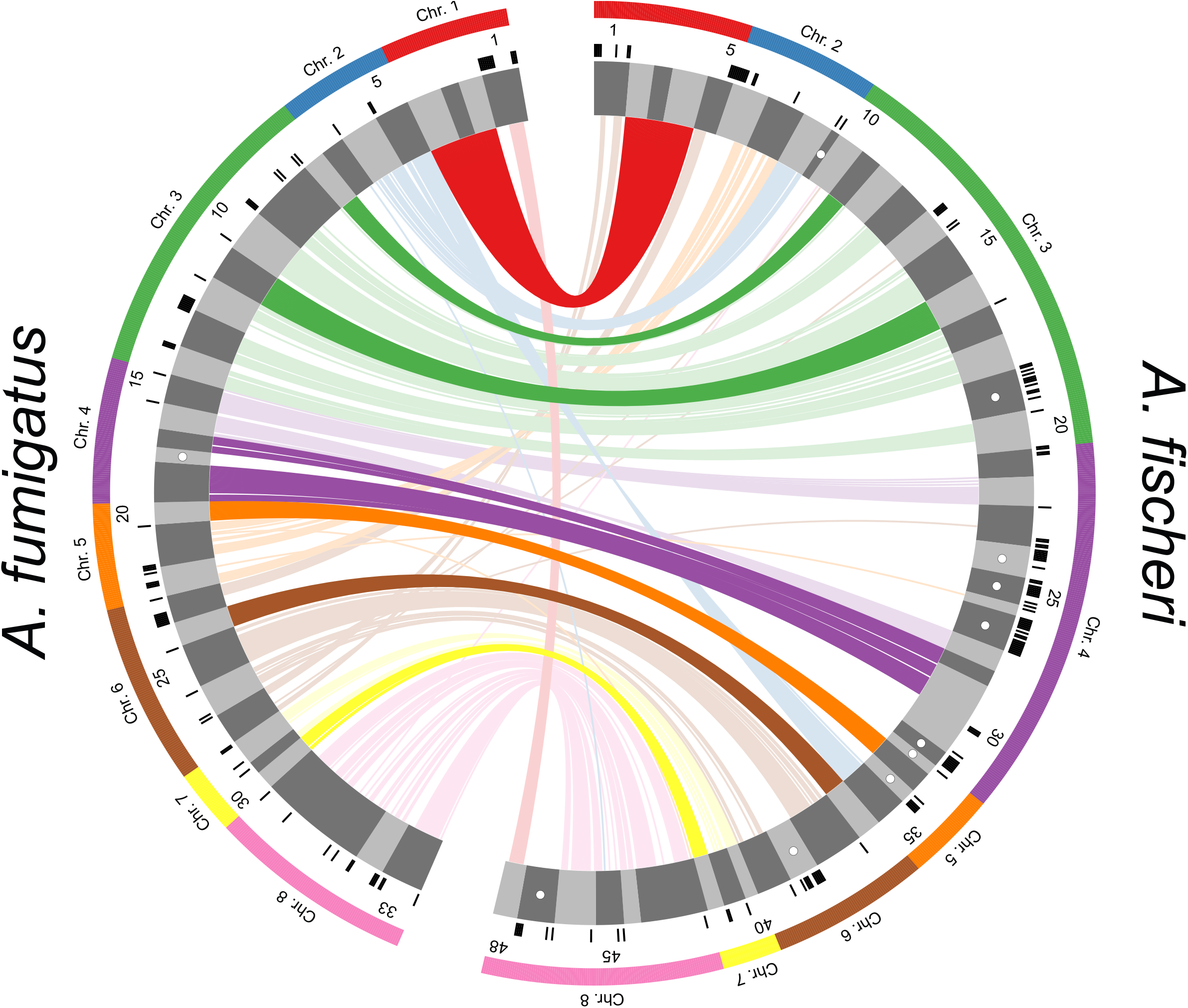
Secondary Metabolite Clusters of *A. fumigatus* and *A. fischeri* show substantial evolutionary divergence. Predicted secondary metabolite gene clusters are shown in the inner track, are alternatively colored dark and light gray, and their size is proportional to the number of genes in them. Black ticks on the exterior of the cluster track indicate a gene that possesses an ortholog in the other species but is not in a secondary metabolite gene cluster in the second species. White dots indicate species-specific clusters. Solid bars on the exterior correspond to the chromosome on which the clusters below them reside. Genes are connected to their orthologs in the other species with dark lines if >90% of the cluster genes in *A. fumigatus* are conserved in the same cluster in *A. fischeri*. Lighter lines connect all other orthologs that are present in both species’ sets of secondary metabolite clusters. Image was made using Circos version 0.69-4 (81).

Only one gene cluster (Cluster 18) was completely *A. fumigatus*-specific. Interestingly, our previous examination of the genomes of 66 *A. fumigatus* strains showed that this cluster was a “jumping cluster”, as it was found to be present in only 5 strains and to reside in three distinct genomic locations (40). Conversely, there are 10 *A. fischeri*-specific gene clusters that do not have orthologs in secondary metabolic gene clusters in *A. fumigatus*. One of these gene clusters is responsible for making helvolic acid [a gene cluster known to be absent from the *A. fumigatus* strain CEA10 but present in strain Af293 (40)], but the other 9 have not been biochemically connected to any metabolite.

All the genes required for the production of the mycotoxin gliotoxin are located in a gene cluster in *A. fischeri* (Fig. S5), and are in fact similar to their *A. fumigatus* orthologs (41), even though *A. fischeri* is not known to produce this mycotoxin (42). Both the gliotoxin and acetylaszonalenin gene clusters are adjacent to one another in the *A. fischeri* genome (Fig. S5). In *A. fumigatus*, the gliotoxin gene cluster is immediately next to what appears to be a truncated version of the acetylaszonalenin cluster that lacks portions of the nonribosomal peptide synthase and acetyltransferase genes as well as the entire indole prenyltransferase gene required for acetylaszonalenin production. The close proximity of these two gene clusters is noteworthy, as it is similar to previously reported “super clusters” in *A. fumigatus* and *A. fumigatus*-related strains (43). These super clusters have been hypothesized to be “evolutionary laboratories” that may give rise to new compounds and pathways (40).

### Isolation and characterization of three new compounds from *A. fischeri*

The relatively low level of conservation of secondary metabolic gene clusters we observed between *A. fumigatus* and *A. fischeri* led us to characterize the secondary metabolites produced by *A. fischeri* (Fig. S6) (44-48). The one strain-many compounds (OSMAC) approach was used to alter the secondary metabolites being biosynthesized in order to produce a diverse set of molecules (49-52). Depending on the media on which it was grown, *A. fischeri* produced as few as 4 (Yeast Extract Soy Peptone Dextrose Agar - YESD) or as many as 10 compounds (Oatmeal Agar – OMA) (Fig. S7). These results showed that culture media influences the biosynthesis of secondary metabolites in *A. fischeri*, a phenomenon observed in many other fungi (50, 53).

To characterize the peaks of interest we observed when *A. fischeri* was grown on OMA, we increased the size of our fungal cultures; doing so yielded seven previously isolated compounds (sartorypyrone A (**1)**, aszonalenin (**4**), acetylaszonalenin (**5**), fumitremorgin A (**6**), fumitremorgin B (**7**), verruculogen (**8**), and the C-11 epimer of verruculogen TR2 (**9**)) and three newly biosynthesized secondary metabolites (sartorypyrone E (**2**), 14-epi-aszonapyrone A (**3**), and 13-*O*-fumitremorgin B (**10**). Two of the secondary metabolites were new compounds (**2** and and one was a new natural product (**10**)) (Fig. 5B). The structures for all 10 compounds were determined using a set of spectroscopic (1 and 2D NMR) and spectrometric techniques (HRMS). Our data for sartorypyrone A (**1**) (54), aszonalenin (**4**) (55, 56), acetylaszonalenin (**5**) (54, 57), fumitremorgin A (**6**) (58, 59), fumitremorgin B (**7**) (60-62), verruculogen (**8**) (63, 64), and the C-11 epimer of verruculogen TR2 (**9**) (64) correlated well with literature values. The structures of 14-epi-aszonapyrone A (**3**), and 13-*O*-prenyl fumitremorgin B (**10**) were fully characterized in this study (see Figshare document: https://doi.org/10.6084/m9.figshare.7149167); the structure elucidation of sartorypyrone E (**2**) is ongoing and will be reported in detail in a forthcoming manuscript.

**Figure 5:**
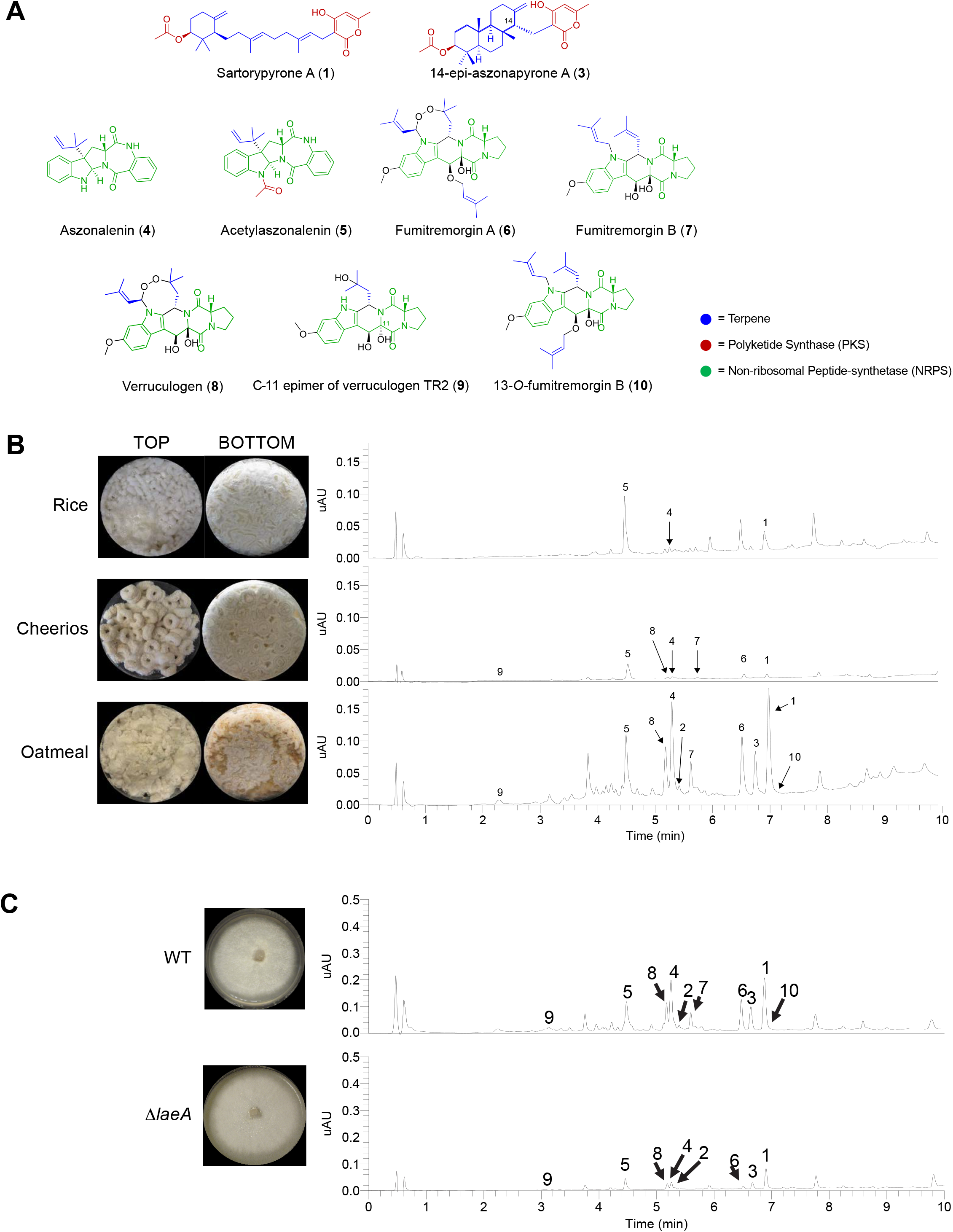
Secondary metabolite production in *A. fischeri*. A) Compounds isolated from *A. fischeri*: (1) sartorypyrone A, (2) sartorypyrone E, (3) 14-epimer aszonapyrone A, (4) aszonalenin, (5) acetylaszonalenin, (6) fumitremorgin A, (7) fumitremorgin B, (8) verruculogen, (9) C-11 epimer verruculogen TR2, and (10) 13-*O*-prenyl-fumitremorgin B. The color coding indicates which putative class the molecule belongs to; e.g., terpenes, PKS, or NRPS. B) Top, *Aspergillus fischeri* was initially grown on rice for two weeks, and then extracted using methods outlined in Fig. S6. The rice culture yielded compounds **1**, **4**, and **5**. Middle, *A. fischeri* was grown on multigrain Cheerios for two weeks, which yielded compounds **1** and **4**-**9**. Bottom, *A. fischeri* on Quaker oatmeal for two weeks. All compounds that were previously isolated in rice and multigrain cheerios cultures in addition to three new compounds (**2**, **3**, and **10**) were found in the oatmeal culture. All pictures depict fungi growing in 250 mL Erlenmeyer flasks; left panel indicates top view, while the right panel shows bottom view. All chromatographic profiles have been normalized to the highest μAU value. C) *Aspergillus fischeri* WT and Δ*laeA* were grown on solid breakfast oatmeal for two weeks and extracted using organic solvents as indicated previously. The crude de-sugared and de-fatted extracts were run using UPLC-MS at a concentration of 2 mg/mL with 5 μL being injected for analysis. The chromatographic profiles were normalized to the highest μAU value. Mass spec analysis indicated the presence of secondary metabolites **1**–**10** within the wild type, and only **1-6**, **8**, and **9** were seen in the Δ*laeA* mutant. All pictures show *A. fischeri* grown on oatmeal agar in Petri plates.

Since four secondary metabolites (**5**-**8**) from *A. fischeri* had also been reported from *A. fumigatus*, we hypothesized that the mechanisms *A. fischeri* employs to regulate its secondary metabolism would also be similar to those used by *A. fumigatus*. To test this hypothesis, we constructed a deletion mutant of *laeA* in *A. fischeri* (Fig. S8). LaeA is a master regulator of secondary metabolism in *A. fumigatus* and a variety of other fungi (65-67). Both the wild type and Δ*laeA* strains of *A. fischeri* were subjected to LC-MS analysis. The chromatographic profile of Δ*laeA* showed mass data that corresponded to sartorypyrone A (**1**), sartorypyrone E (**2**), 14-epi-aszonapyrone A (**3**), aszonalenin (**4**), acetylaszonalenin (**5**), fumitremorgin A (**6**), verruculogen (**8**), and the C-11 epimer of verruculogen TR2 (**9**). However, the relative abundance of compounds present was very low compared to the wild type (Fig. 5C). Fumitremorgin B (**7**) and 13-*O*-prenyl-fumitremorgin B (**10**) were not produced by the Δ*laeA* mutant at all.

## Discussion

*A. fumigatus* is a major human fungal pathogen, yet its close relative *A. fischeri* is rarely an agent of human disease. A number of traits that contribute to the virulence of *A. fumigatus* have been characterized, but their distribution and potential role in *A. fischeri-*mediated disease was largely unknown. In this study, we thoroughly characterized *A. fischeri* (strain NRRL 181) and compared it to *A. fumigatus* (strain CEA10) for multiple disease-relevant biological and chemical differences. Our data shows that *A. fischeri* can grow in a mammalian host but is much less fit and causes a disease progression quite different than that observed during *A. fumigatus* infections (Figs. 1 and 2). Further investigations revealed that secondary metabolic genes are much less conserved than genes in the rest of the genome (Fig. S4), and a chemical analysis of *A. fischeri* resulted in the identification of both previously identified and new compounds (Fig. 5). While the biosynthetic pathways producing secondary metabolites in *A. fischeri* and *A. fumigatus* appear to be quite different, our data suggest that a master regulator of secondary metabolism in *A. fumigatus* (*laeA*) possesses a similar role in *A. fischeri* (Fig. 5C).

In order to cause disease, a microbe must be able to respond to the set of diverse and stressful environments presented by its host. Based on our data, *A. fischeri* is unable to respond to many of these stresses as well as *A. fumigatus* (Figs. 2-3). We hypothesize that this inability to thrive under stress contributes to the varying disease progressions observed during our animal model experiments (Fig. 1). Some or all of the genetic determinants responsible for this discrepancy in stress response and virulence could reside in the 9 *A. fumigatus*-specific genes we identified (Fig. S4); alternatively, some of the ~1,700 *A. fischeri*-specific genes we identified may inadvertently facilitate control of *A. fischeri* in a mammalian host. Importantly, our analyses herein were conducted on single strains of the two species and further studies are needed to determine how representative these observed trait and genomic differences are across strains.

Even though more than 10% of the genes in each species lack an ortholog in the other species, only ~2% (1/49) of previously identified genetic determinants of virulence in *A. fumigatus* are not conserved in *A. fischeri* (Table S1). This result, and our observation that many of the pathways of secondary metabolism are quite different between *A. fischeri* and *A. fumigatus*, support a multifactorial model of *A. fumigatus* virulence (1, 68, 69) and suggest a need to investigate virulence on multiple levels of biological complexity. In order to cause disease in a host, *A. fumigatus* (and other species closely related to it) must adhere and germinate in the lung (69), survive inherently stressful conditions presented by host environments (ex. severe lack of metals and oxygen) (6, 70, 71), and modulate or endure actions of the host immune system (72). Given the diversity of these activities, it is unlikely that single genes or pathways will be responsible for the totality of *A. fumigatus*-derived disease, even though not all genes in the genome have been characterized for their role in pathogenicity. We hypothesize that multiple pathways (including those involved in secondary metabolism) have changed during the evolution of *A. fischeri* and *A. fumigatus*, resulting in their differing ability to cause disease.

*A. fumigatus* and *A. fischeri* are members of *Aspergillus* section *Fumigati*, a clade that includes multiple closely related species, some of which are pathogens (e.g., *A. fumigatus*, *A. lentulus*, and *A. udagawae*) and some of which are considered non-pathogens (e.g., *A. fischeri*, *A. aureolus*, and *A. turcosus*) (2, 42, 73, 74). The ability to cause disease in humans appears to have either arisen or been lost (or both) multiple times independently during the evolution of this lineage, as pathogenic species are spread throughout the phylogeny (17, 75). A broader, phylogenetically-informed comparison of pathogenic and non-pathogenic species in section *Fumigati* would provide far greater resolution in identifying (or dismissing) factors and pathways that may contribute or prevent the ability of these organisms to cause disease. Also, leveraging the diversity of section *Fumigati* would give researchers a better understanding of the nature and evolution of human fungal pathogenesis as the appreciation for the health burden caused by fungi increases (76).

An important caveat to our experiments is that we only analyzed a single, representative strain from each species. Several recent studies have identified a wide variety of differences between *A. fumigatus* strains, which have in turn been shown to contribute to physiological differences, including but not limited to secondary metabolism and virulence (30, 40, 72, 73). While the genome of only one isolate of *A. fischeri* has so far been sequenced (16) and the organism has only been reported to cause human disease a few times (21-24), it would be of great interest to compare patient-derived and environment-derived isolates at the genomic, phenotypic, and chemical levels. Although it appears that clinical and environmental isolates do not stem from separate lineages in *A. fumigatus* (77), whether this is also the case for largely non-pathogenic species, such as *A. fischeri*, or for rarely isolated pathogenic species, such as *A. lentulus* or *A. udagawae*, remains largely unknown.

## Materials and Methods

### Strains and growth media

*A. fischeri* strain NRRL 181 was acquired from the ARS Culture Collection (NRRL). *A. fumigatus* strain CEA10 (CBS 144.89) was obtained from the CBS. All strains were grown on glucose minimal media (GMM) from conidial glycerol stocks stored at −80°C. All strains were grown in the presence of white light at 37°C. Conidia were collected in 0.01% Tween-80 and enumerated with a hemocytometer.

### Murine virulence studies

For the chemotherapeutic (leukopenic) murine model, outbred CD-1 female mice (Charles River Laboratories, Raleigh, NC, USA), 6-8 weeks old, were immunosuppressed with intraperitoneal (i.p.) injections of 150 mg/kg cyclophosphamide (Baxter Healthcare Corporation, Deerfield, IL, USA) 48 hours before and 72 hours after fungal inoculation, along with subcutaneous (s.c.) injections of 40 mg/kg Kenalog-10 (triamcinolone acetonide, Bristol-Myer Squibb, Princeton, NJ, USA) 24 hours before and 6 days after fungal inoculation. For the murine triamcinolone model outbred CD-1 female mice, 6-8 weeks old, were treated with 40 mg/kg Kenalog-10 by s.c. injection 24 hours prior to fungal inoculation.

For both models, conidial suspensions of 2 × 10^6^ conidia were prepared in 40 μL sterile PBS and administered to mice intranasally while under isoflourine anesthesia. Mock mice were given 40 μL PBS. Mice were monitored three times a day for signs of disease for 14 or 18 days post-inoculation. Survival was plotted on Kaplan-Meir curves and statistical significance between curves was determined using Mantel-Cox Log-Rank and Gehan Breslow-Wilcoxon tests. Mice were housed in autoclaved cages at 4 mice per cage with HEPA filtered air and autoclaved food and water available at libitum.

### *Galleria mellonella* virulence studies

*G. mellonella* larvae were obtained by breeding adult moths (78). *G. mellonella* larvae of a similar size were selected (approximately 275–330 mg) and kept without food in glass container (Petri dishes), at 37°C, in darkness for 24 h prior to use. *A. fumigatus* and *A. fischeri* conidia were obtained by growing on YAG media culture for 2 days. The conidia were harvested in PBS and filtered through a Miracloth (Calbiochem). The concentration of conidia was estimated by using hemocytometer, and resuspended at a concentration of 2.0 × 10^8^ conidia/ml. The viability of the conidia was determined by incubating on YAG media culture, at 37°C, 48 hours. Inoculum (5 μl) of conidia from both strains were used to investigate the virulence of *A. fumigatus* and *A. fischeri* against *G. mellonella*. Ten *G. mellonella* in the final (sixth) instar larval stage of development were used per condition in all assays. The control group was the larvae inoculated with 5 μl of PBS to observe the killing due to physical trauma. The inoculum was performed by using Hamilton syringe (7000.5KH) and 5 μl into the haemocel of each larva via the last left proleg. After, the larvae were incubated in glass container (Petri dishes) at 37°C in the dark. The larval killing was scored daily. Larvae were considered dead by presenting the absence of movement in response to touch.

### Histopathology

Outbred CD-1 mice, 6-8 weeks old, were immunosuppressed and intranasally inoculated with 2 × 10^6^ conidia as described above for the chemotherapeutic and corticosteroid murine models. Mice were sacrificed 72 hours post inoculation. Lungs were perfused with 10% buffered formalin phosphate before removal, then stored in 10% buffered formalin phosphate until embedding. Paraffin embedded sections were stained with haematoxylin and eosin (H&E) and Gömöri methenamine silver (GMS). Slides were analyzed microscopically with a Zeiss Axioplan 2 imaging microscope (Carl Zeiss Microimaging, Inc. Thornwood, NY, USA) fitted with a Qimiging RETIGA-SRV Fast 1394 RGB camera. Analysis was performed in Phylum Live 4 imaging software.

### Ethics Statement

We carried out our mouse studies in strict accordance with the recommendations in the Guide for the Care and Use of Laboratory Animals of the National Research Council (Council, 1996). The mouse experimental protocol was approved by the Institutional Animal Care and Use Committee (IACUC) at Dartmouth College (Federal-Wide Assurance Number: A3259-01).

### Growth Assays

Radial growth was quantified by point inoculation of 1 × 10^3^ conidia in 2 μL on indicated media; plates were incubated at 37°C in normoxia (~21% O_2_, 5% CO_2_) or hypoxia (0.2% O_2_, 5% CO_2_). Colony diameter was measured every 24 hours for 4 days and reported as the average of three biological replicates per strain.

For 2-DG experiments, 1 × 10^3^ conidia in 2 μL were spotted on 1% lactate minimal media with or without 0.1% 2-deoxyglucose (2-DG; Sigma, D8375). Plates were incubated for 3 days at 37°C in normoxia or hypoxia with 5% CO_2_. Percent inhibition was calculated by dividing radial growth on 2-DG plates by the average radial growth of biological triplicates on plates without 2-DG.

Fungal biomass was quantified by measuring the dry weight of fungal tissue from 5 × 10^7^ conidia grown in 100 mL liquid GMM shaking at 200 rpm for 48 hours in normoxia (~21% O_2_) and hypoxia (0.2% O_2_, 5% CO_2_). Liquid biomass is reported as the average of three biological replicates per strain. Hypoxic conditions were maintained using an INVIVO_2_ 400 Hypoxia Workstation (Ruskinn Technology Limited, Bridgend, UK) with a gas regulator and 94.8% N_2_.

Liquid growth curves were performed with conidia adjusted to 2 × 10^4^ conidia in 20 μL 0.01% Tween-80 in 96-well dishes, then 180 μL of media (GMM or lung homogenate) was added to each well. Plates were incubated at 37°C for 7 hours, then Abs_405_ measurements were taken every 10 minutes for the first 16 hours of growth with continued incubation at 37°C. Lung homogenate media was prepared as follows: lungs were harvested from healthy CD-1 female mice (20-24 g) and homogenized through a 100 μM cell strainer in 2 mL PBS/lung. Homogenate was diluted 1:4 in sterile PBS, spun down to remove cells, then filter sterilized through 22 μM PVDF filters.

### Cell wall and oxidative stresses

Congo Red (0.5 mg/mL), Menadione (20 μM), or calcofluor white (CFW, 25 μg/mL) were added to GMM plates. 1 × 10^3^ conidia (Calcofluor white and Menadione) or 1 × 10^5^ conidia (Congo Red) were point inoculated and plates were incubated for 96 hours at 37°C with 5% CO_2_.

### Orthology Determination and Analyses

Genomes for *A. fumigatus* CEA10 and *A. fischeri* NRRL 181 were downloaded from NCBI (Accession numbers of GCA_000150145.1 and GCF_000149645.1, respectively). To identify putative orthologous genes between *A. fischeri* and *A. fumigatus*, a reciprocal best BLAST hit (RBBH) approach was used. We blasted the proteome of *A. fischeri* to *A. fumigatus* and vice versa using an e-value cutoff of 10^−3^ and then filtered for RBBHs according to bitscore (79). A pair of genes from each species was considered orthologous if their best blast hit was to each other. Species-specific and orthologous protein sets were visualized using version 3.0.0 of eulerAPE (80).

### Secondary Metabolism Cluster Prediction and Analyses

Version 4.2.0 of antiSMASH (38) was used with its default settings to identify secondary metabolite clusters. Orthologous cluster genes were identified using our RBBH results and visualized using version 0.69 of Circos (81). Chromosomes were identified for *A. fischeri* NRRL1 and *A. fumigatus* CEA10 using NUCMER (82)and chromosomal sequences from *A. fumigatus* strain AF293 from NCBI (Accession number GCA_000002655.1). Syntenic clusters were visualized using easyfig version 2.2.2 (83).

### Secondary Metabolite Extraction and Identification

Secondary metabolites were extracted from *A. fischeri* using techniques well established in the Natural Products literature (84, 85). This was done by adding a 1:1 mixture of CHCl_3_:CH_3_OH and left to shake overnight. The resulting slurry was partitioned twice, first with a 4:1:5 CHCl_3_:CH_3_OH:H_2_O solution, with the organic layer drawn off and evaporated to dryness *in vaccuo*, and secondly reconstituting 1:1:2 CH_3_CN:CH_3_OH:hexanes, where the organic layer was drawn off and evaporated to dryness. The extract then underwent chromatographic separation (flash chromatography and HPLC) using varied gradient systems. The full structural characterization of the new secondary metabolites is provided in the Figshare document (https://doi.org/10.6084/m9.figshare.7149167).

### Construction of the *A. fischeri* Δ*laeA* mutant

The gene replacement cassettes were constructed by *‘‘in vivo’’* recombination in *S. cerevisiae* as previously described by (86, 87). Approximately 2.0 kb from the 5’-UTR and 3’-UTR flanking regions of the targeted ORF regions were selected for primer design. The primers pRS NF010750 5’fw (5’-GTAACGCCAGGGTTTTCCCAGTCACGACGCAGTCTAACGCTGGGCCCTTCC-3’) and pRS NF010750 3’rv (5’-GCGGTTAACAATTTCTCTCTGGAAACAGCTACGGCGTTTGACGGCACAC-3’) contained a short homologous sequence to the Multicloning site (MCS) of the plasmid pRS426. Both the 5’-and 3’-UTR fragments were PCR-amplified from *A. fischeri* genomic DNA (gDNA). The *prtA* gene, conferring resistance to pyrithiamine, which was placed within the cassette as a dominant marker, was amplified from the pPRT1 plasmid by using the primers prtA NF010750 5’rv (5’-GTAATCAATTGCCCGTCTGTCAGATCCAGGTCGAGGAGGTCCAATCGG-3’) and prtA NF010750 3’fw (5’-CGGCTCATCGTCACCCCATGATAGCCGAGATCAATCTTGCATCC-3’). The deletion cassette was generated by transforming each fragment along with the plasmid pRS426 cut with *Bam*HI/*Eco*RI into the *S. cerevisiae* strain SC94721, using the lithium acetate method (88). The DNA from the transformants was extracted by the method described by Goldman et al. (89). The cassette was PCR-amplified from these plasmids utilizing TaKaRa Ex Taq™ DNA Polymerase (Clontech Takara Bio) and used for *A. fisheri* transformation according to the protocol described by Malavazi and Goldman (87). Southern blot and PCR analyses were used to demonstrate that the cassette had integrated homologously at the targeted *A*. *fischeri* locus. Genomic DNA from *A. fischeri* was extracted by grinding frozen mycelia in liquid nitrogen and then gDNA was extracted as previously described (87). Standard techniques for manipulation of DNA were carried out as described (90). For Southern blot analysis, restricted chromosomal DNA fragments were separated on 1% agarose gel and blotted onto Hybond N^+^ nylon membranes (GE Healthcare). Probes were labeled using [α-^32^P]dCTP using the Random Primers DNA Labeling System (Life Technologies). Labeled membranes were exposed to X-ray films, which were scanned for image processing. Southern blot and PCR schemes are shown in Fig. S8.

## Acknowledgements

Computational infrastructure was provided by The Advanced Computing Center for Research and Education (ACCRE) at Vanderbilt University. MEM, JS, and AR were supported by a Vanderbilt University Discovery Grant. Research in AR’s lab is also supported by the National Science Foundation (DEB-1442113), the Guggenheim Foundation, and the Burroughs Wellcome Fund. RAC holds an Investigator in the Pathogenesis of Infectious Diseases Award supported by the Burroughs Wellcome Fund and is also supported by a National Institute of Allergy and Infectious Diseases award 1R01AI130128. SRB was supported, in part, by the National Institute of General Medical Sciences of the National Institutes of Health under Award Number T32GM008704. SLK was supported by the National Center for Complementary and Integrative Health, a component of the National Institutes of Health, under award number T32 AT008938. GHG was supported by grants from Fundação de Amparo à Pesquisa do Estado de São Paulo (FAPESP) and Conselho Nacional de Desenvolvimento Científico e Tecnológico (CNPq), both from Brazil.

## Supplementary Material

**Figure S1: *A. fumigatus* grows slower than *A. fischeri* in glucose minimal media (GMM), but at the same speed as *A. fischeri* in lung homogenate media.** *A. fumigatus* CEA10 or *A. fischeri* NRRL181 were cultured in flat-bottom 96 well plates at 2 × 10^4^ conidia per well. Conidia were added in a 20 μL of 0.01% Tween-80 and media was carefully pipetted over the inoculum into each well. Lung homogenate was generated according to (29). Plates were incubated for 7 hours at 37°C before measurements at 405 nm were taken every 10 min. Mean and SEM of eight technical replicates; data is representative of three biological replicates.

**Figure S2: *A. fischeri* and *A. fumigatus* exhibit similar growth patterns at 30**°**C.** 1 × 10^3^ conidia were point inoculated on each plate then plates were incubated at 30°C in normoxia (~21% oxygen, 5%CO_2_); colony diameter was measured every 24 hours. Mean and SEM of triplicates. Tween-80 – 1% Tween-80 provided as sole carbon source; CAA – Casamino acids; GMM – glucose minimal media.

**Figure S3: In contrast to *A. fumigatus*, *A. fischeri* fails to thrive at 44**°**C.** Error bars indicate standard deviations between biological duplicates (**P-value < 0.005 in a paired, equal variance student t-test).

**Figure S4: The genomes of *A. fumigatus* and *A. fischeri* are largely similar, but their secondary metabolic pathways are quite divergent.** Left, Venn diagram showing the sets of *A. fischeri*-specific proteins, shared orthologous proteins, and *A. fumigatus*-specific proteins encoded in each genome. Numbers below each species name indicate the total number of proteins encoded in that genome. Right, Venn diagram showing the sets of *A. fischeri*-specific secondary metabolite cluster proteins, shared secondary metabolite cluster genes, and *A. fumigatus*-specific secondary metabolite cluster genes. Numbers below each species name indicate the total number of secondary metabolite cluster proteins encoded in that genome. In each diagram, circles are proportional to the number of proteins they contain.

**Figure S5: The acetylaszonalenin and gliotoxin clusters in *A. fumigatus* and *A. fischeri* are located immediately next to one another.** The portions of Clusters 37 and 25 from *A. fischeri* and *A. fumigatus*, respectively, that are known to contain the previously characterized acetylaszonalenin (91) and gliotoxin (41) clusters is shown. Genes colored in shades of green are involved in the acetylaszonalenin biosynthetic pathway. Dark green, *anaPS* (nonribosomal peptide synthase). Light green, *anaAT* (acetyltransferase). Green, *anaPT* (prenyltransferase). Orange, gliotoxin biosynthetic genes. Gray arrow, syntenic gene in both species not involved in gliotoxin synthesis. Sequences that are similar to one another (based on blastn scores) are marked by gray parallelograms. Image was made using EasyFig version 2.2.2 (83).

**Figure S6: A custom chemical analysis protocol was developed for studying the metabolites produced by *A. fischeri*.** Approximately 60 mL of 1:1 CH_3_OH:CH_3_Cl was added to cultures of *Aspergillus fischeri* grown on solid-state fermentation for two weeks. The cultures were then chopped thoroughly with a large scalpel and shaken for 16 hours using an orbital shaker. The liquid culture was then vacuum filtered and concentrated using 90 mL CH_3_Cl and 150 mL water and transferred into a separatory funnel. The organic (bottom) layer was drawn off and evaporated to dryness. The dried, de-sugared extract was reconstituted in 100 mL of 1:1 CH_3_OH:CH_3_CN and 100 mL of hexane. The biphasic solution was shaken vigorously and transferred to a separatory funnel. The CH_3_OH:CH_3_CN layer was evaporated to dryness under vacuum, producing a de-fatted extract. The extract was then subdivided into several peaks or fractions using flash chromatography. The subfractions were further separated using HPLC until pure compounds were isolated. The pure compounds were subjected to UPLC-MS analysis to establish the molecular formula and fragmentation patterns. Finally, pure compounds were identified using both NMR analysis as well as information from UPLC-MS data.

**Figure S7: *A. fischeri* produces different numbers of metabolites, depending on the media it is grown on.** Base peak chromatograms as measured by LC-MS, illustrating how the chemistry profiles varied based on growth conditions. PDA + ab was used as the chemical control to observe the differences in the secondary metabolites, due to it being the media that *A. fischeri* is stored. There were overall no chemical differences observed between the different variations of PDA media. Each peak (which indicates different chemical entities) was observed in the three PDA variations, albeit at fluctuating intensities. SDA, PYG, and YESD produced the majority of the peaks observed in PDA, but it also lacked some observed peaks, indicating that these growth conditions were not chemically favored. CYA produced the majority of the peaks, as well as an additional peak that was observed at a much lower intensity in PDA. However, this peak was similarly observed in OMA. OMA produced similar peaks to those observed in PDA, but with higher intensity. Due to this, OMA was selected to further study. The gray boxes indicate differences in the observed peaks compared to PDA. See Figshare document (https://doi.org/10.6084/m9.figshare.7149167) for more information.

**Figure S8: Southern blot confirms construction of the** Δ***laeA* mutant.** A 1kb probe recognizes a single DNA band (~4.4kb) in the wild type strain and a single DNA band (~2.7kb) in the Δ*laeA* mutant.

All supplemental tables can be found on Figshare (https://doi.org/10.6084/m9.figshare.7149167)

**Table S1: Virulence-associated genes in *A. fumigatus* and *A. fischeri*.**

**Table S2: Bioinformatically predicted secondary metabolite clusters in *A. fumigatus* strain CEA10.**

**Table S3: Bioinformatically predicted secondary metabolite clusters in *A. fischeri* strain NRRL 181.**

**Table S4: Different Types of Growth Media used for *Aspergillus fischeri*.**

